# Phytochemical profiling and antioxidant potential of freshwater algal extracts from Lahore, Pakistan, with preliminary evaluation of cytotoxic activity

**DOI:** 10.64898/2026.05.11.724325

**Authors:** Saman Safdar Rehan, Aasia Kiran, Ghazal Yasmeen, Awaise Altaf, Tahir Maqbool, Faheem Hadi, Saira Aftab

## Abstract

Freshwater algae represent an underexplored source of naturally occurring bioactive metabolites with potential applications in pharmaceutical and biomedical research. This study investigated the phytochemical composition, antioxidant capacity, and preliminary cytotoxic potential of ethanolic and n-hexane extracts of freshwater algal species collected at Jilani Park, Lahore, Pakistan. Algal species were identified morphologically by Dr. Ghazal Yasmeen (Institute of Botany, Punjab University, Lahore). Extracts were analyzed using gas chromatography–mass spectrometry (GC–MS) and qualitative phytochemical screening. Antioxidant activity was evaluated using DPPH radical scavenging, hydrogen peroxide scavenging, and reducing power assays. Cytotoxic potential was assessed using MTT and cell adhesion assays on HeLa and SF767 cell lines as preliminary indicators of bioactivity.

GC–MS analysis identified 25 compounds, including sterols, fatty acid esters, terpenoids, phenolic compounds, and volatile metabolites. Phytochemical screening confirmed the presence of flavonoids, phenolics, tannins, and terpenoids in the extracts. Among the tested extracts, the n-hexane fraction demonstrated comparatively higher antioxidant activity across multiple assays. Ethanolic extracts showed moderate reductions in HeLa cell viability, whereas limited effects were observed in SF767 cells.

These findings suggest that freshwater algae are promising natural reservoirs of antioxidant metabolites with potential relevance for future isolation and characterization of bioactive compounds for biomedical applications. Further purification and mechanistic studies are required to identify specific active constituents.

## Introduction

In recent years, advanced extraction approaches, including supercritical fluid extraction, have increasingly been applied for the efficient isolation of bioactive compounds from natural sources, allowing improved recovery of biologically active metabolites with potential biomedical relevance (Fabrowska et al., 2016).

A wide range of secondary metabolites contribute to biological defense systems in algae and other photosynthetic organisms (De *et al*., 2000). Several microalgal species have been investigated as natural sources of functional metabolites with antioxidant and other biologically relevant properties (Rodriguez *et al*., 2008; Najdenski *et al*., 2013; Abbas *et al*., 2016). These metabolites include tannins, phenolic acids, lignins, alkaloids, flavonoids, coumarins, and terpenoids, which are widely distributed in plant and algal tissues and contribute to ecological adaptation and chemical defense mechanisms (Ahmad *et al*., 2011). Many of these compounds have attracted attention due to their antioxidant potential and possible future applications in compound isolation and pharmaceutical research.

Microalgae comprise a diverse group of simple photosynthetic organisms that may be unicellular or filamentous and are highly efficient in converting solar energy into biologically active biomass. Filamentous green macroalgae are particularly abundant in freshwater ecosystems and are increasingly recognized as environmentally and biotechnologically valuable resources. Their cultivation has also been explored for applications in nutrient assimilation, bioremediation, and removal of organic pollutants from aquatic systems (Henkanatte *et al*., 2015; Kang and Wen, 2015; Kumar *et al*., 2015).

*Cladophora glomerata* is a widely distributed freshwater macroalga found in nutrient-rich aquatic environments throughout the world (Higgins *et al*., 2008). It is characterized by a branched filamentous structure and commonly occurs in eutrophic lakes and freshwater habitats (John *et al*., 2015). Previous studies have shown that C. glomerata contains diverse phytochemicals including flavonoids, phenolics, fatty acids, tannins, sterols, alkaloids, and terpenoids (Mohamed *et al*., 2013). These metabolites have been associated with antioxidant and other biological activities, highlighting the species as a potentially valuable source of naturally occurring bioactive compounds for future extraction and characterization studies (Laungsuwon *et al*., 2013; Awad *et al*., 2009; Surayot *et al*., 2016).

## Introduction

Species of the green algal genus *Rhizoclonium (Cladophorales*) are distributed across marine, brackish, and freshwater habitats, although they are more frequently reported from brackish ecosystems (John *et al*., 2002; Leliaert *et al*., 2016). *Rhizoclonium kützingii* is characterized by thin, unbranched filaments composed of cylindrical cells, some of which produce rhizoids. Molecular investigations based on nuclear-encoded small subunit ribosomal DNA sequences have suggested phylogenetic complexity within this genus, indicating possible polyphyletic relationships (Hanyuda *et al*., 2002).

Another important freshwater species, *Rhizoclonium hieroglyphicum*, is recognized as an edible alga with a polysaccharide-rich cell wall and seasonal growth patterns influenced by environmental conditions such as temperature and water flow. Extracts of this species contain structurally diverse biomolecules, including polysaccharides composed of arabinose, xylose, rhamnose, and galactose, as well as amino acids such as phenylalanine, tyrosine, and tryptophan (Mungmai *et al*., 2014). These constituents suggest that freshwater *Rhizoclonium* species may represent promising natural reservoirs of antioxidant and structurally valuable metabolites for future biochemical and pharmacological investigation.

Cancer remains a major global health challenge and continues to be a leading cause of mortality worldwide (Rennie & Rusting, 1996). Despite considerable advances in treatment strategies, including chemotherapy and combination-based therapeutic approaches, adverse side effects and variable clinical responses continue to limit treatment efficacy (Varshney *et al*., 2013). Consequently, there is sustained interest in identifying naturally derived bioactive compounds with diverse biological properties that may support future pharmaceutical investigation.

Natural products have historically provided structurally diverse metabolites with broad biological activities, including antioxidant and cytotoxic effects. Several algae-derived compounds have demonstrated the ability to modulate cellular processes associated with growth regulation, including inhibition of proliferation, induction of cell cycle arrest, regulation of oxidative stress pathways, and modulation of programmed cell death mechanisms (Xiao *et al*., 2007; Lee *et al*., 2004; Tzianabos, 2000). These observations have encouraged continued exploration of freshwater algal species as potential sources of biologically active metabolites for future isolation and mechanistic characterization.

Evaluation of crude algal extracts using established human cell lines provides an effective preliminary approach for identifying biologically active fractions. In the present study, HeLa and SF767 cell lines were employed as representative in vitro models to assess the cytotoxic potential of freshwater algal extracts and to provide an initial indication of cellular bioactivity.

SF767 is a human glioblastoma-derived cell line frequently used as an experimental model for studying cellular responses in glioma-associated systems. Glioblastomas are aggressive central nervous system tumors derived from glial cells and are classified by the World Health Organization as grade IV astrocytomas due to their highly proliferative nature and poor clinical prognosis (Louis *et al*., 2007). These tumors remain associated with limited survival despite multimodal treatment approaches including surgery, radiotherapy, and chemotherapy (Jakubowicz *et al*., 2010). Glioblastoma belongs to a broader group of glial-derived neoplasms that include astrocytomas, oligodendrogliomas, and ependymomas (Zhang *et al*., 2012).

The use of both HeLa and SF767 cell lines in this study enabled comparative evaluation of crude extract-associated cytotoxicity across distinct cellular backgrounds, allowing preliminary assessment of extract bioactivity while supporting future studies aimed at identifying specific antioxidant-associated metabolites responsible for these effects.

## Materials and Methods

### Collection and Identification of Algal Material

Freshwater algal samples were collected in September 2020 from the pond located in Jilani Park, Lahore, Pakistan. Samples were collected aseptically in sterile polyethylene bags, transferred to labeled plastic containers, and transported to the laboratory for processing.

A small portion of each sample was mounted on clean glass slides and examined microscopically for preliminary morphological identification. Additional portions were preserved in 4% formalin for detailed taxonomic examination. Species identification was performed by Dr. Ghazal Yasmeen using standard taxonomic keys and morphological characteristics.

Samples were thoroughly washed with distilled water to remove debris and epiphytes, air-dried at room temperature away from direct sunlight, ground into fine powder, and stored at 18°C until extraction.

### Preparation of Algal Extracts

Dried algal powder (50 g) was separately extracted with 500 mL of n-hexane and absolute ethanol in conical flasks. The mixtures were incubated at room temperature on an orbital shaker for 72 h to facilitate solvent penetration and metabolite extraction. Extracts were filtered through Whatman No. 1 filter paper, and solvents were removed under reduced pressure using a rotary evaporator to obtain concentrated crude extracts as previously described (Rasool *et al*., 2014 and Kalsoom *et al*., 2024).

The crude extracts were stored at -20°C until further analysis.

### Cell Culture Maintenance

Human cervical epithelial (HeLa) and glioblastoma-derived (SF767) cell lines were obtained from the School of Biological Sciences (Punjab University) and maintained in DMEM high-glucose medium supplemented with 10% fetal bovine serum/FBS ((Sigma-AldrichLot#BCBS3184V) and 1% penicillin–streptomycin solution (Caisson Lot#10201011)).

Cells were incubated at 37°C under 5% CO_2_ and 95% relative humidity and passaged every 2–3 days using 0.25% trypsin–EDTA. For assays, almost 20 thousand cells per well of a 96-well plate were seeded out for 24h, and then crude extracts were exposed in 2% FBS containing medium for 48 h.

### Morphological Assessment

Cell morphology following treatment was observed under Floid® Imaging station for bright field setting and 20x magnification.

### MTT Cytotoxicity Assay

Metabolic activity of cells under treatment (100 and 200 ug/ml) was evaluated using the MTT assay, and for that purpose, cells were seeded into 96-well plates and allowed to attach overnight before treatment with different extract concentrations for 48h. Following incubation, MTT reagent was added, and plates were incubated for 2 h in the MTT working solution. The supernatant was removed, and formazan crystals were dissolved in DMSO. Optical density was measured at 570 nm. (Life Technology® recommended protocol)

Cell viability was calculated as:

Cell viability (%) = (Absorbance of treated cells / Absorbance of control cells) × 100

### Cell Adhesion Assay

Cell adhesion was assessed by seeding HeLa and SF767 cells in 96-well plates, followed by treatment with algal extracts (100 and 200 μg/mL). Cells were fixed and stained with 0.1% crystal violet. After washing with PBS, the dye was solubilized using Tween solution and absorbance was recorded at 570 nm (Lukas *et al*., 2019; Aftab *et al*, 2019).

### Qualitative Phytochemical Screening

Qualitative phytochemical screening was performed according to standard protocols described by Savithramma *et al*. (2011) to determine the presence or absence of major secondary metabolites including alkaloids, steroids, tannins, terpenoids, saponins, phenols, flavonoids, coumarins, quinones, and glycosides.

### Determination of Total Phenolic Content

Total phenolic content was determined using the Folin–Ciocalteu reagent method with minor modifications (Terpinc et al., 2012). Absorbance was measured at 750 nm using a Shimadzu double-beam UV–Visible spectrophotometer (Canan et al., 2016). Results were expressed as mg gallic acid equivalent (GAE)/100 mg dry extract.

### Determination of Total Flavonoid Content

Total flavonoid content was determined according to the method described by Chang et al. (2002). Absorbance was recorded at 415 nm using a Shimadzu UV–Visible spectrophotometer (Gupta, 2015).

### Determination of Total Tannin Content

Total tannin content was quantified using Folin–Denis reagent and expressed as tannic acid equivalent (TAE).

### DPPH Radical Scavenging Assay

Antioxidant activity was evaluated using the DPPH free radical scavenging assay as described by Koleva and Van Beek *et al*. (2002). Reduction in absorbance indicated increased radical scavenging activity and was calculated as:

% Scavenging activity = [1 − (Absorbance of sample / Absorbance of control)] × 100

### Reducing Power Assay

Reducing power was determined based on the reduction of ferricyanide (Fe^3+^) to ferrocyanide (Fe^2+^), followed by the formation of ferric–ferrous complex. Absorbance was measured at 700 nm, with ascorbic acid used as a standard reference.

### Hydrogen Peroxide Scavenging Assay

Hydrogen peroxide scavenging activity was evaluated according to Gulçin et al. (2004). Absorbance was measured at 230 nm, and scavenging activity was calculated as: % Scavenging = [(A_0_ − A_1_)/A_0_] × 100

where A_0_ represents control absorbance, and A_1_ represents sample absorbance.

### GC–MS Analysis

Phytochemical profiling was performed using gas chromatography–mass spectrometry (GC–MS) at H.E.J. Research Institute of Chemistry.

Analysis was conducted using a Shimadzu GCMS-QP2010 Ultra system equipped with an Rxi-5Sil MS capillary column (30 m × 0.25 mm × 0.25 μm). Helium was used as carrier gas at a constant flow rate of 1.0 mL/min. The injector temperature was maintained at 250°C. The oven temperature program was initiated at 70°C and increased to 280°C at a rate of 10°C/min. Electron ionization was performed at 70 eV over a scan range of m/z 40–600.

Compounds were identified by comparison of retention times and mass spectral fragmentation patterns with entries in the NIST mass spectral library.

### Molecular Docking

Molecular docking analysis was performed using AutoDock Vina to evaluate the binding affinity of selected algal-derived compounds against the Akt target protein. The docked protein–ligand complexes were prepared and visualized using PyMOL. Interactions between ligands and amino acid residues within the active binding site were analyzed using PLIP. Two-dimensional and three-dimensional interaction images were generated using Discovery Studio Client for structural interpretation. The drug-likeness properties of the top-ranked compounds were predicted using the online SwissADME platform, while pharmacokinetic and toxicity profiles were estimated using pk CSM

## Results

### Morphological Observations

Morphological observations were made using a bright-field setting on the Imaging station by Life Technology^™^

In the case of HeLa cell lines the morphological observation showed that the untreated and DMSO treated samples show no difference as compared to the control samples while ethanolic extract treated 100µg/ml samples had lower cell density and cell shape was bit elongated. The n-Hexane treatment showed no difference in cell number or cell morphology.

In the case of SF767, both untreated and DMSO had no significant difference but the ethanolic extract treated cells had significant difference in 200µg/ml concentrations as compared to the 100µg/ml concentration with cell density very low and cell shape round to the oval. In n-Hexane treated cells no difference in the cell morphology or cell density was observed in any of the concentration.

### Cytotoxic Screening of HeLa and SF767 Cells

Morphological examination revealed concentration-dependent alterations following ethanolic extract treatment, particularly in HeLa cells, including reduced cell density and mild cellular elongation at higher concentrations. Minimal morphological changes were observed in n-hexane-treated cells.

In SF767 cells, ethanolic extract treatment at 200 μg/mL resulted in reduced cell density and altered cellular morphology, whereas n-hexane treatment showed limited observable effects.

MTT analysis demonstrated dose-dependent reduction in metabolic activity following ethanolic extract exposure in HeLa cells, with significant reduction at 200 μg/mL. No statistically significant reduction was observed in SF767 cells.

Cell adhesion analysis indicated reduced adherence in HeLa cells treated with ethanolic extract (100 μg/mL). In SF767 cells, both ethanolic and n-hexane extracts showed moderate effects at selected concentrations.

These findings indicate measurable but modest cytotoxic bioactivity of crude freshwater algal extracts and support further fractionation studies to identify specific metabolites responsible for these effects.

### Phytochemical and anti oxidant analysis by biochemical assays

The phytochemical screening was performed with ethanol, hexane, and chloroform of freshwater algae. Glycosides did not show any positive result for their presence in any of the four extracts tested, as shown in Table 1.

**Table 1.** The quantitative analysis of phytochemical substances in algal extracts.

|  | **Total Flavonoid Content** | **Total Phenol Content** | **Total Tannin Content** |
| --- | --- | --- | --- |
| **Ethanol** | 70.01266±4.617 | 3.5189±0.299 | 30±1.310 |
| **Hexane** | 7.521315±3.563 | 3.59528±0.042 | 30.0389±2.131 |

Among the three different extracts, hexane showed the presence of the maximum number of compounds. Ethanolic extracts showed six compounds, and chloroform only five compounds. Ethanolic extract showed the presence of steroids, tannins, terpenoids, saponins, phenols, and flavonoids. Hexane extract showed the presence of tannins, terpenoids, saponins, phenols, flavonoids, coumarins, and quinones. The presence or absence of the phytochemicals depends upon the solvent medium used for extraction.

The highest total flavonoid contents were observed in ethanolic extract (70.01266±4.617) which was higher than other extracts. It was observed that chloroform extract have lowest flavonoid content (3.5023±1.924). The highest phenolic content were observed in ethanol and hexane extracts (3.5189±0.299 and 3.59528±0.042) respectively.

Higher tannin content observed in hexane extract (30.0389±2.131) as compared to ethanolic extract. The ethanolic extract has lower content (30±1.310) as shown in table 1.

The effect of antioxidants on DPPH radical is thought to be due to their hydrogen donating ability. The extracts of fresh water algal samples allowed reacting with stable DPPH. The reduction capability of DPPH radical is determined at 515nm absorbance. The hexane extract showed highest free radical scavenging activity (55.4702±3.354) compared to the ethanolic extract (26.1898±4.031). The free radical scavenging potential of the freshwater algal compounds was analyzed according to (Muller et al., 2011) with slight modifications.

The table 3 indicated that hexane have highest content (75.2±0.338) as compared to ethanolic extract (69.5±2.460).

The hexane has higher fatty acid content 124.444±0.880 as compared to ethanolic extract 41.994±0.946.

Hexane extracts showed higher content (71.14286±0.874) as compared to ethanolic extract 38.7619±0.464).

### GC–MS Profiling

GC–MS analysis identified 25 compounds in freshwater algal ethanolic crude extract based on preliminary data collected from cytotoxic studies on human cell lines. Detected metabolites included fatty acid esters, sterols, volatile compounds, and saturated and unsaturated hydrocarbons. γ-Sitosterol and vitamin E were among the identified bioactive constituents. Several unidentified peaks were also observed, suggesting the presence of structurally novel metabolites requiring further characterization.

### Molecular Docking

Among the screened compounds, 24,25-Dihydroxycholecalciferol exhibited the highest docking score against Akt; however, in silico toxicity prediction suggested potential hepatotoxicity. Therefore, cholesterol, which demonstrated the second-highest docking affinity and showed comparatively favorable predicted pharmacokinetic characteristics, was also considered for further interpretation. Depending on the intended biological emphasis of the study, either compound may be discussed, although cholesterol may provide a more conservative and pharmacologically relevant candidate if exclusion of potentially hepatotoxic ligands is preferred.

Descriptions and interpretation criteria for PKCSM-derived pharmacokinetic parameters were based on previously published studies employing comparable computational prediction workflows.

## Discussion

The phytochemical composition of freshwater algal extracts was found to be strongly influenced by the polarity of the extraction solvent, which determined the distribution and detectability of secondary metabolites. Screening of ethanol, hexane, and chloroform extracts revealed the presence of several major classes of bioactive compounds, including flavonoids, phenolics, tannins, terpenoids, steroids, coumarins, and quinones, whereas glycosides were not detected in any of the tested extracts (Tables 1 and 2). These findings are consistent with previous reports demonstrating that solvent polarity plays a critical role in extracting specific classes of phytoconstituents from algal biomass.

**Table 2:** Antioxidant activities.

| Sr. No. | DPPH | H<sub>2</sub>O<sub>2</sub> | RPA | FA |
| --- | --- | --- | --- | --- |
| **Ethanol** | $26.1898 \pm 4.031$ | $38.7619 \pm 0.464$ | $69.5 \pm 2.460$ | $41.994 \pm 0.946$ |
| **Hexane** | $55.4702 \pm 3.354$ | $71.14286 \pm 0.874$ | $75.2 \pm 0.338$ | $124.444 \pm 0.880$ |

**Table 3:** Comparative summary of phyto chemical Analysis.

| **Sr. No.** | **Phytochemical Parameters** | **Ethanol** | **Hexane** |
| --- | --- | --- | --- |
| **1.** | **Steroids** | + | - |
| **2.** | **Tannins** | + | + |
| **3.** | **Terpenoids** | ++ | + |
| **4.** | **Saponins** | + | + |
| **5.** | **Phenols** | + | ++ |
| **6.** | **Flavonoids** | + | + |
| **7.** | **Coumarins** | - | + |
| **8.** | **Quinones** | - | + |
| **9.** | **Glycosides** | - | - |

Among the detected metabolites, tannins were consistently present across all extracts. Tannins are polyphenolic compounds widely reported for their biological relevance, including antioxidant, antimicrobial, and anti-inflammatory properties (Kolodziej & Kiderlen, 2005). Their presence in natural extracts has also been associated with inhibitory effects against a range of pathogenic microorganisms, including Escherichia coli, Staphylococcus aureus, and Salmonella typhi (Scalbert, 1991; Chung et al., 1998). In addition, tannin-rich preparations have been investigated for their potential role in oxidative stress modulation and related biological processes (Cowan, 1999; Heil et al., 2002).

Terpenoids were detected in ethanol and hexane extracts. These compounds are widely recognized for their structural diversity and biological activities, including antioxidant and cytotoxic properties in various natural systems (Abid et al., 1997; Valls et al., 1995; Culioli et al., 2001). Similarly, steroids were present in ethanol and chloroform extracts and are known to contribute to a range of biological activities, including membrane stabilization and modulation of enzymatic processes. Naturally occurring steroids have also been reported in nutritional, cosmetic, and medicinal applications due to their broad functional properties (Okwu, 2001).

Flavonoids and phenolic compounds were detected across most extracts, highlighting their widespread distribution in freshwater algae. These metabolites are of particular interest due to their well-documented antioxidant potential and their role in mitigating oxidative stress through radical scavenging mechanisms. Phenolic compounds, in particular, have been associated with a variety of biological activities, including antioxidant, anti-inflammatory, and antimicrobial effects (Aliyu et al., 2009). The observed distribution of phenolic compounds across extracts suggests that freshwater algae may serve as a promising natural reservoir for antioxidant-rich bioactive constituents.

Coumarins were identified only in hexane extracts, whereas quinones were also restricted to the same fraction. Coumarins are known for diverse pharmacological properties, including anticoagulant activity and potential relevance in vascular-related conditions (Jeeva et al., 2012). Quinones, on the other hand, have been reported to exhibit biological activity through interactions with nucleic acids and interference with mitochondrial electron transport processes, often leading to the generation of reactive oxygen species (Solanki et al., 2008). These properties further support the biochemical diversity of freshwater algal metabolites.

The relatively higher phenolic content observed in hexane and ethanol extracts may be attributed to the presence of polyphenolic constituents, which are strongly associated with antioxidant activity. A positive correlation between phenolic content and antioxidant capacity has been widely reported in plant- and algae-derived extracts, where compounds such as flavonoids, tannins, and tocopherols contribute significantly to free radical scavenging (Dapkevicius et al., 1998). In the present study, this relationship was reflected in the enhanced DPPH scavenging activity observed in the hexane fraction, suggesting that non-polar antioxidant constituents may play an important role in radical neutralization.

Antioxidants are defined as compounds capable of delaying or inhibiting oxidative processes by interrupting free radical chain reactions and reducing oxidative damage (Halliwell & Aruoma, 1991). The DPPH assay, a widely used method for evaluating free radical scavenging potential, demonstrated that hexane extracts exhibited the highest antioxidant activity compared to chloroform and ethanol extracts. This suggests that non-polar constituents may contribute significantly to the overall antioxidant capacity of the studied algal species.

Hydrogen peroxide scavenging assays further supported these findings. Hydrogen peroxide, although relatively stable, can generate highly reactive hydroxyl radicals in biological systems, thereby contributing to oxidative stress (Gulçin et al., 2003). The observed concentration-dependent scavenging activity of the algal extracts indicates their potential role in mitigating oxidative species and suggests that these extracts may serve as a source of naturally derived antioxidant compounds.

Morphological evaluation and MTT-based assays were used as preliminary screening tools to assess the cytotoxic potential of freshwater algal extracts against human cell lines. Increasing interest in natural product-derived bioactive compounds has led to extensive screening of algae for potential biological activities. In this study, extracts from Gomphonema incognitum, Cladophora glomerata, Rhizoclonium kützingii, and Rhizoclonium hieroglyphicum demonstrated concentration-dependent effects on cellular morphology and viability.

Ethanolic extracts showed a measurable reduction in cell viability in HeLa cells at higher concentrations, whereas effects in SF767 glioblastoma-derived cells were comparatively limited under similar conditions. These observations should be interpreted as preliminary evidence of crude extract bioactivity rather than specific cytotoxic or therapeutic effects. The variability in response between cell lines further suggests that extract composition and solvent polarity may influence cellular sensitivity.

Overall, the results indicate that freshwater algal extracts contain diverse phytochemicals with notable antioxidant potential and measurable, but non-specific, effects on cell viability in vitro. These findings support the concept that freshwater algae represent a promising natural source of bioactive metabolites that warrant further fractionation, purification, and mechanistic investigation to identify specific antioxidant constituents and their biological roles.

## Statistical analysis

All experiments were performed in triplicate, and data are presented as mean ± standard deviation (SD). Statistical analyses were conducted using GraphPad Prism software. Differences between treated and control groups were evaluated using one-way analysis of variance (ANOVA). A value of p < 0.05 was considered statistically significant with p < 0.05 = *, p < 0.01 = **,p < 0.001 = ***.

## Conclusion

Freshwater algal extracts collected from Lahore contained diverse phytochemical constituents with measurable antioxidant activity and modest cytotoxic effects in preliminary cell-based screening assays. Among the tested extracts, the n-hexane fraction demonstrated comparatively stronger antioxidant performance, suggesting enrichment of non-polar bioactive metabolites.

These findings support the potential of freshwater algae as promising natural sources for the extraction and future characterization of antioxidant compounds of possible biomedical relevance. Further fractionation, purification, and mechanistic studies are required to identify specific active constituents and define their biological properties.

## Implication

The results revealed that extracts contained higher antioxidant activity and consequently, they can be considered as a good source of natural products that may be employed in the treatment of the different diseases that are associated with oxidative stress.

## Data availability statement

Data sharing is not applicable as no new data were generated or analyzed during this study.

## Acknowledgment

We acknowledge the guidance of Dr. Naseema Azeem, Senior Scientist in the School of Biological Sciences, Punjab University, for her help in GC-MS data analysis.

## AI assistance

Tools based on AI were used for language improvement after the main draft had already been developed. No AI tool was used for figure or data modification.

## Figure description

**Figure 1:**
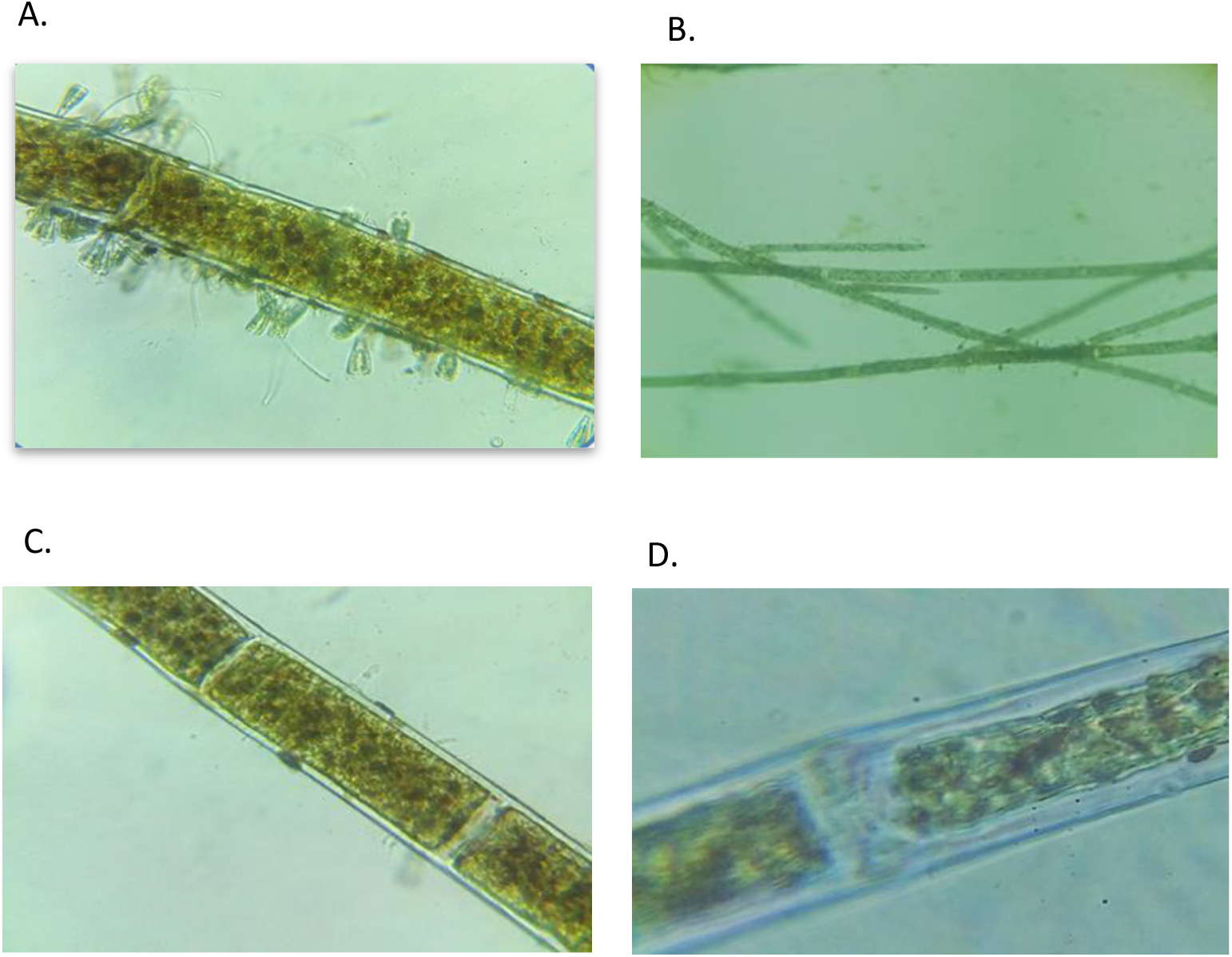
Identification of Algal species: a and b are the microscopic view of *Gomphonema Incognitum &* Microscopic view of *Cladophora Glomerata*. C and d are the microscopic view of *Rhizoclonium Kützing &* Microscopic view of *Rhizoclonium hieroglyphicum* varient.

**Figure 2:**
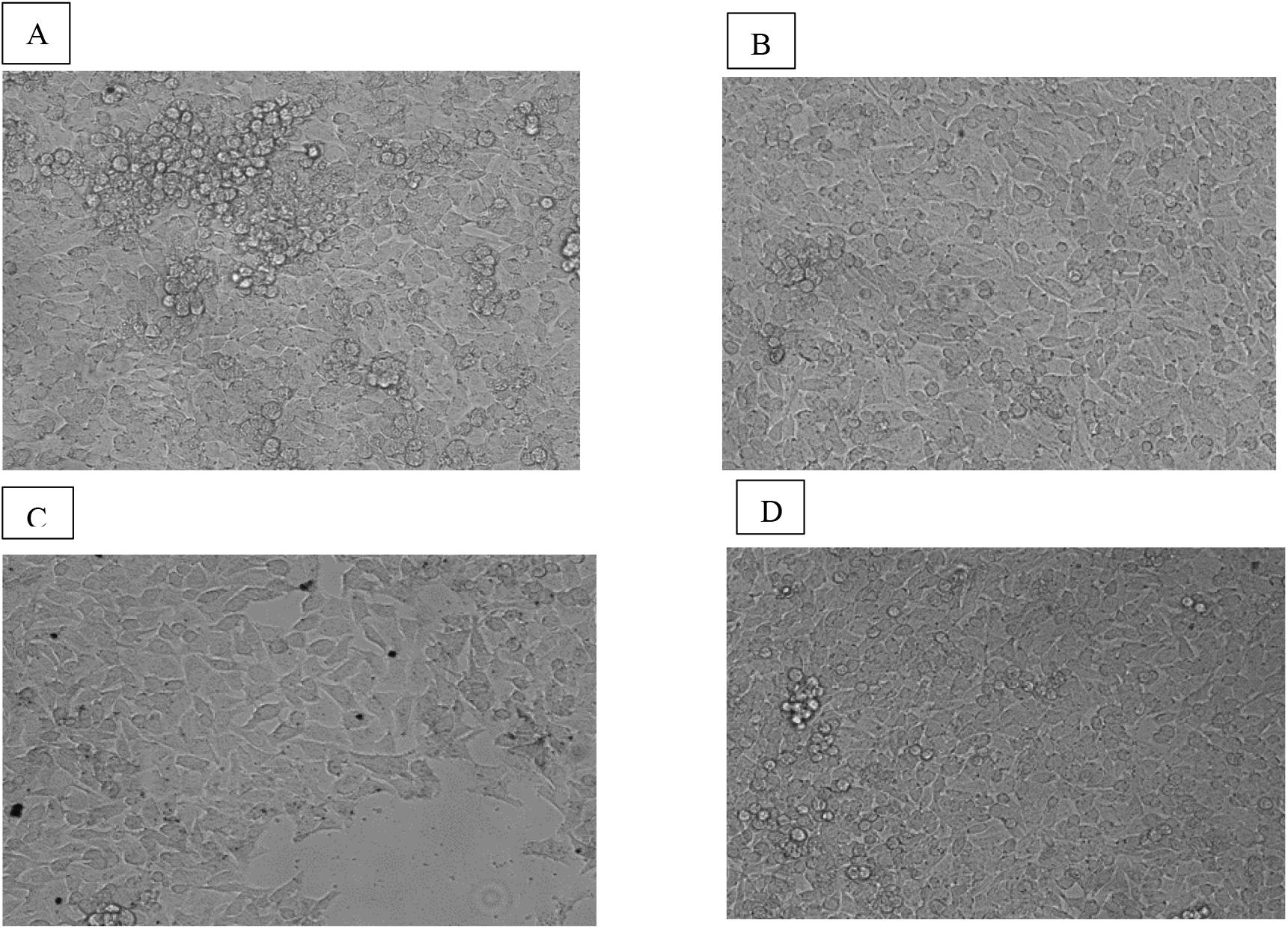
Bright field images of HeLa cell line A) Un-treated B) DMSO treated C) Ethanolic extract treated 100µg/ml and D) n-Hexane extract treated 100µg/ml of freshwater algae.

**Figure 3:**
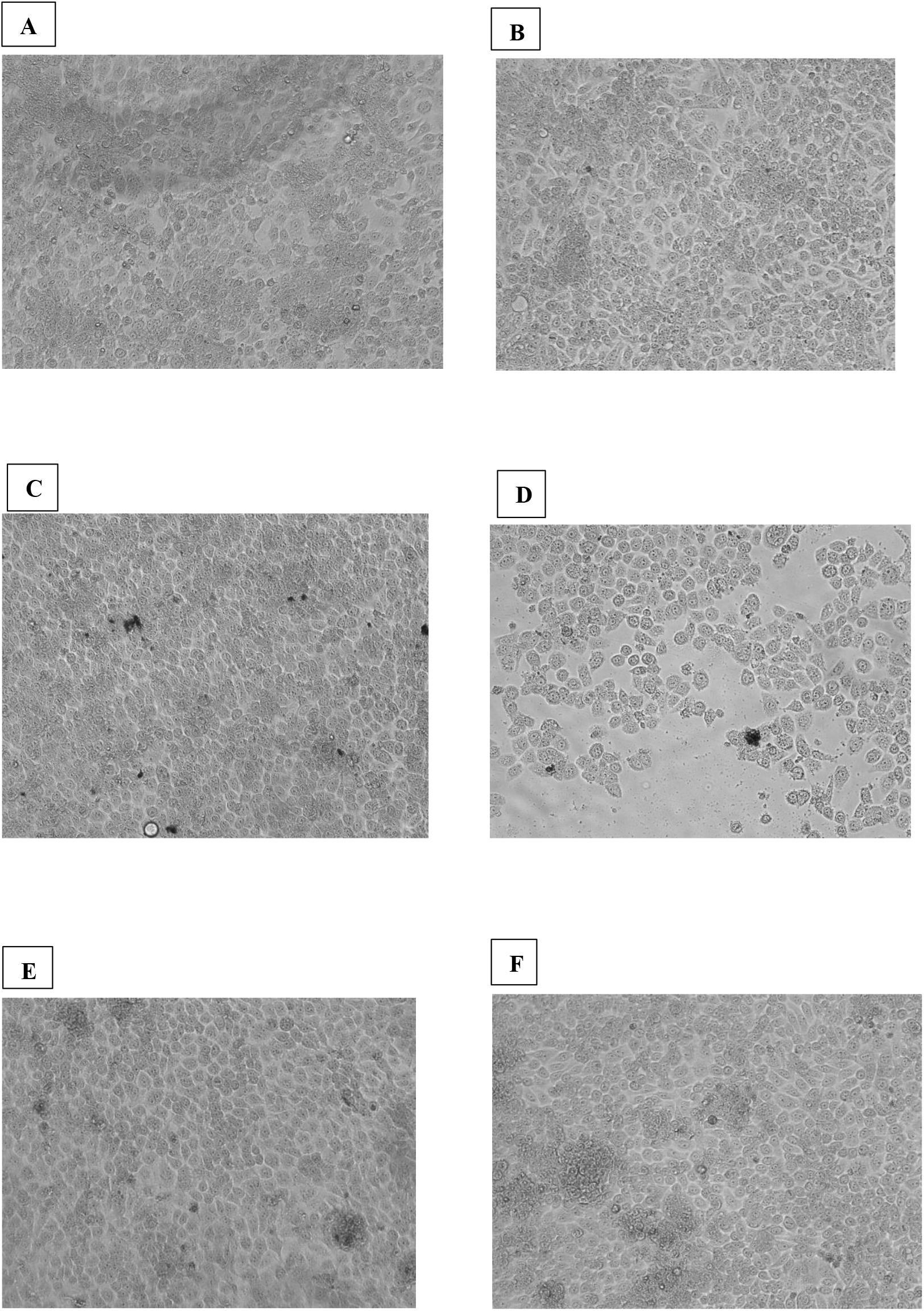
Bright field images of SF767 cell lines. A) Un-treated B) DMSO treated C) Ethanolic extract treated 100µg/ml D) Ethanolic extract treated 200µg/ml, E) n-Hexane extract treated 100µg/ml and F) n-Hexane extract treated 200µg/ml of freshwater algae

**Figure 4:**
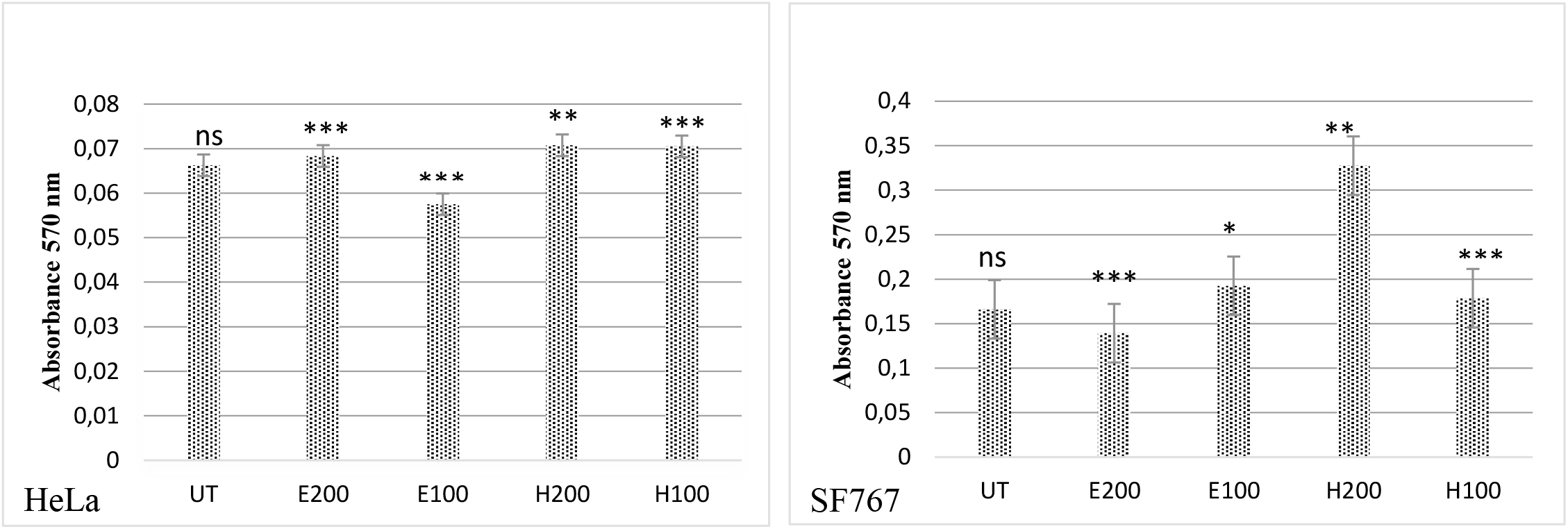
Graph of HeLa cell line and SF767 cell line for Metabolic activity assay using MTT for ethanolic crude extract A. E100 (100µg/ml) E200 (100µg/ml) and Hexanolic crude extract H100 (100µg/ml) and H200 (100µg/ml) normalised with DMSO treated Absorbance as negative control and) compared to Untreated UT samples.

**Figure 5:**
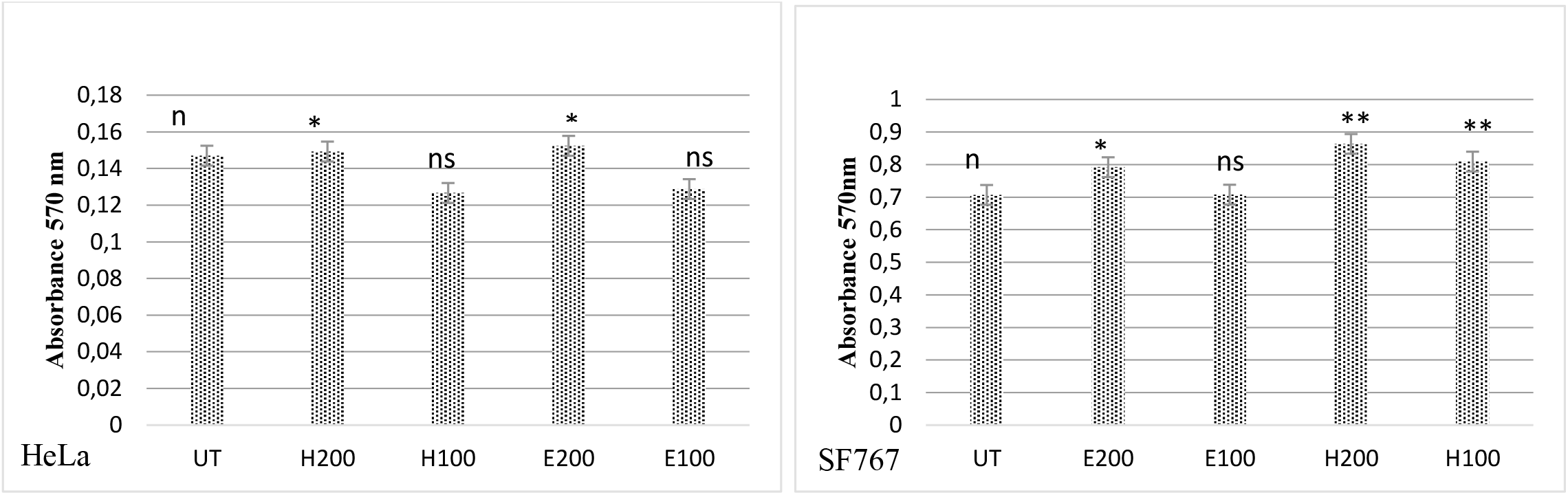
Graph of HeLa cell line and SF767 cell line for cell viability assay using 0.2% Crystal violet for ethanolic crude extract E100 (100µg/ml) E200 (100µg/ml) and Hexanolic crude extract H100 (100µg/ml) and H200 (100µg/ml) normalised with DMSO treated Absorbance as negative control and) compared to Untreated UT samples.

**Figure 6:**
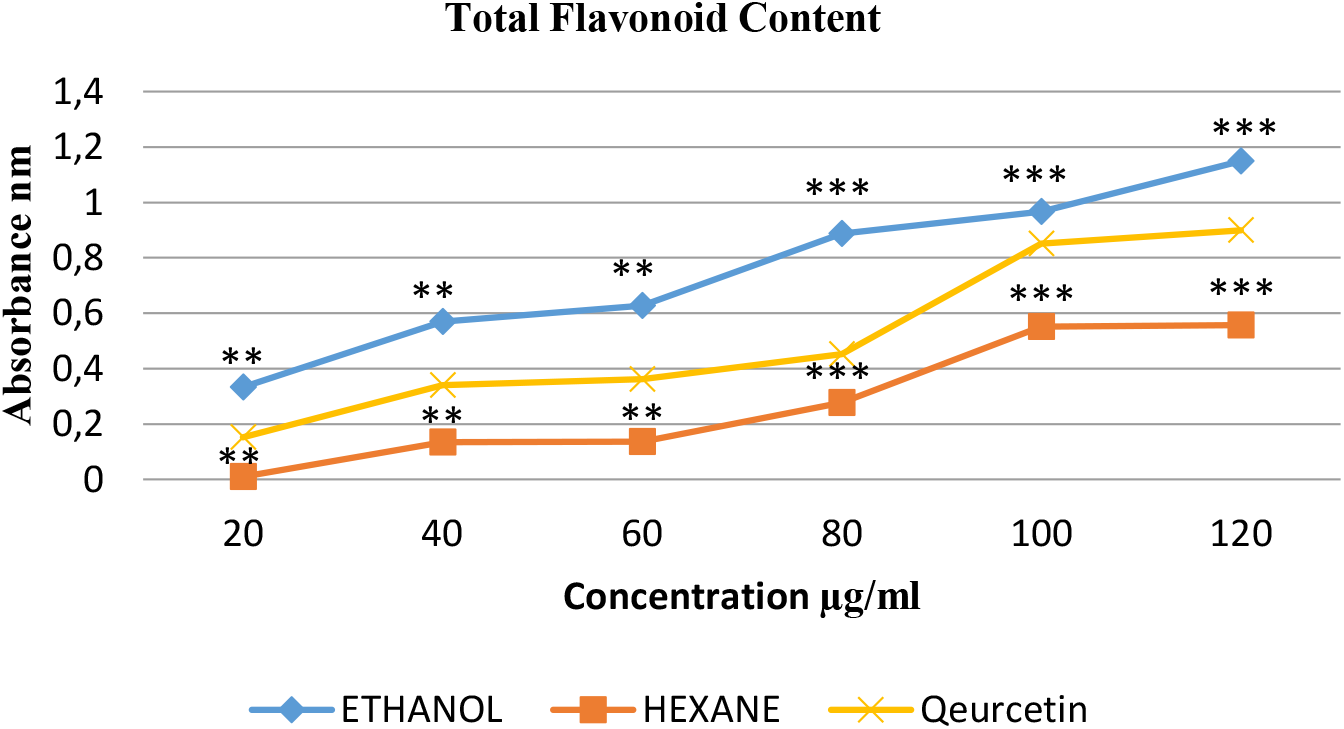
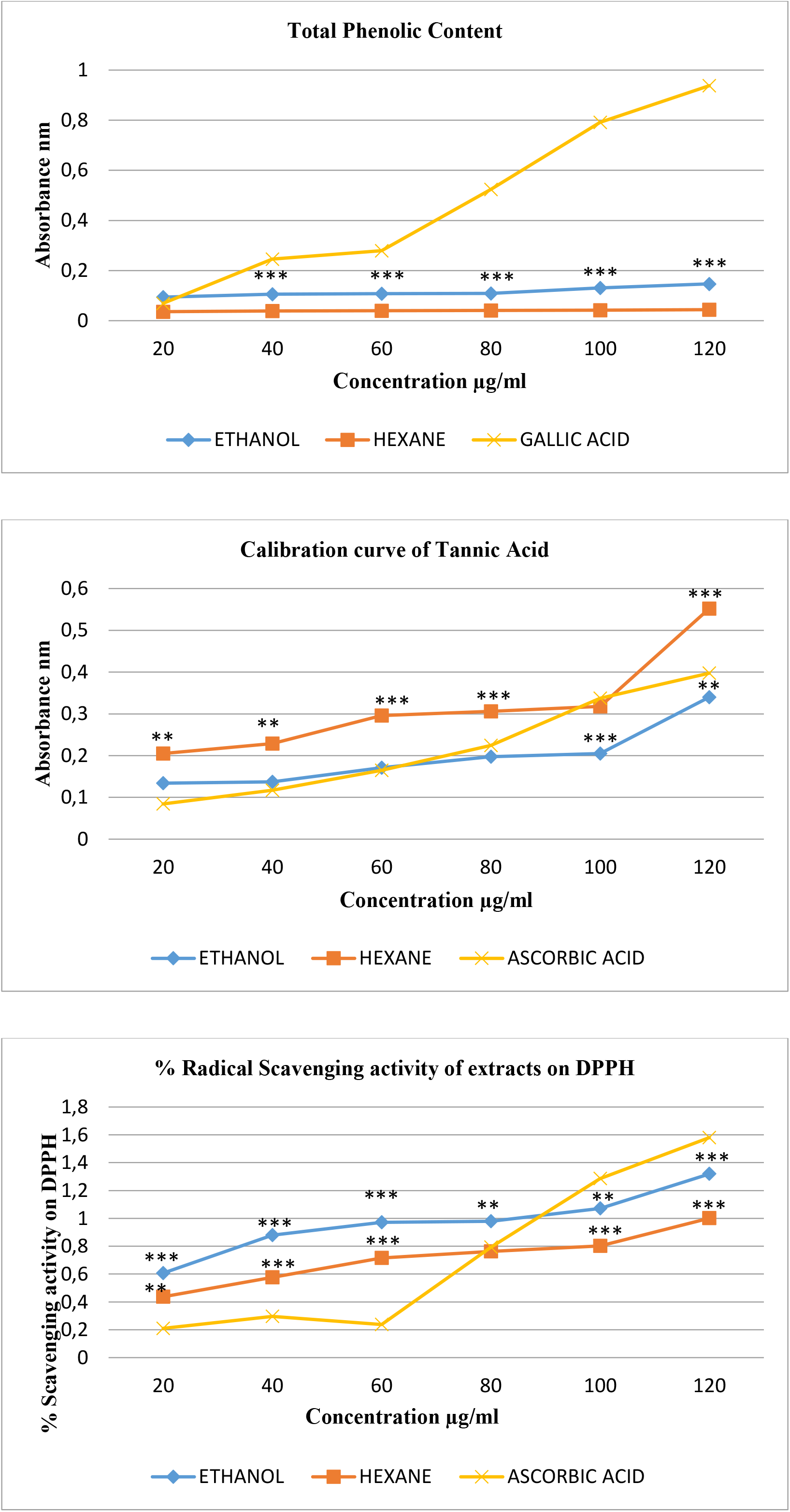
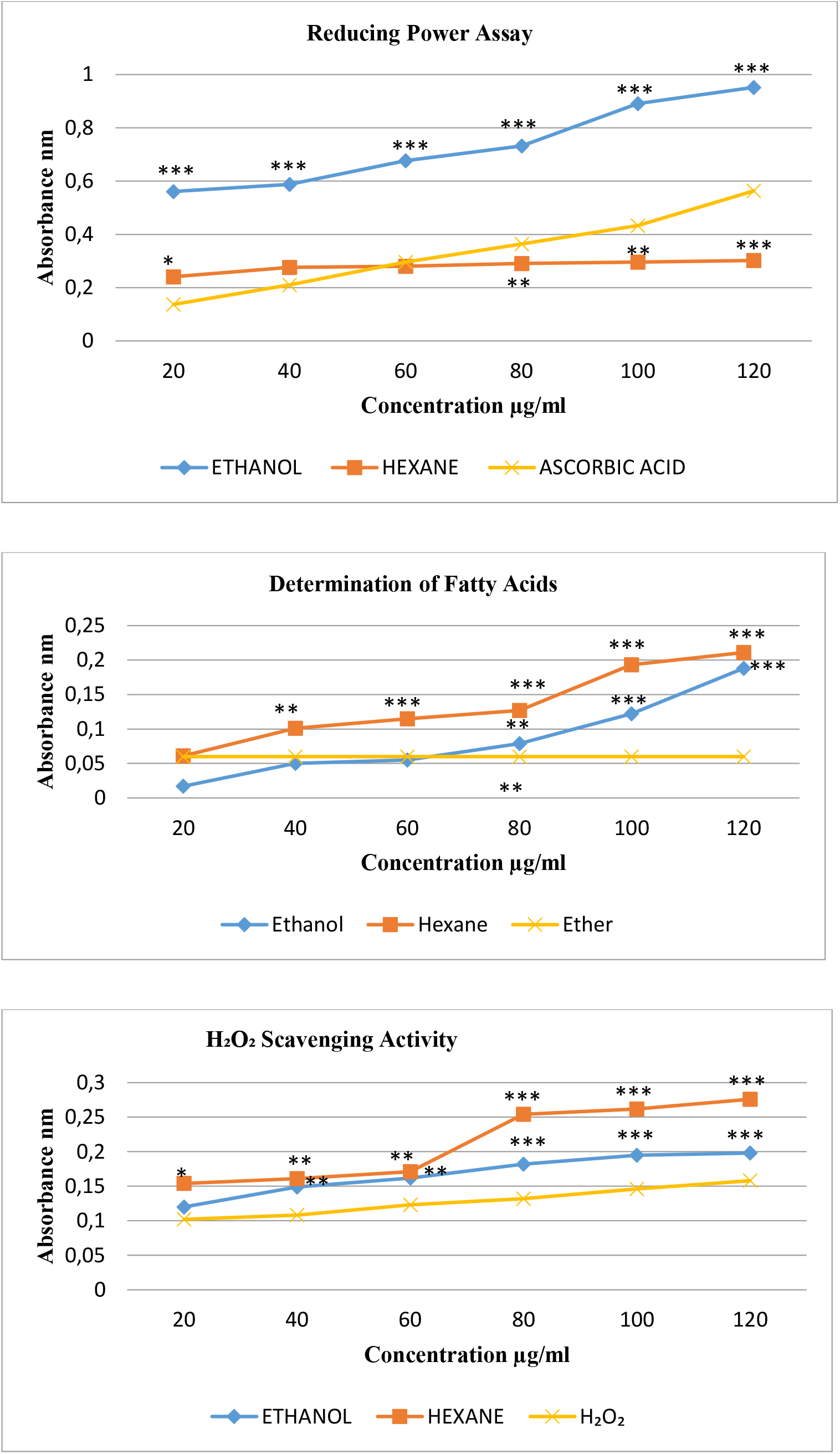
The phytochemical screenings of nine different compounds like steroids, tannins, terpenoids, saponins, phenols, flavonoids, coumarins, quinones, and glycosides along with relevant controls for each compound.

**Figure 7:**
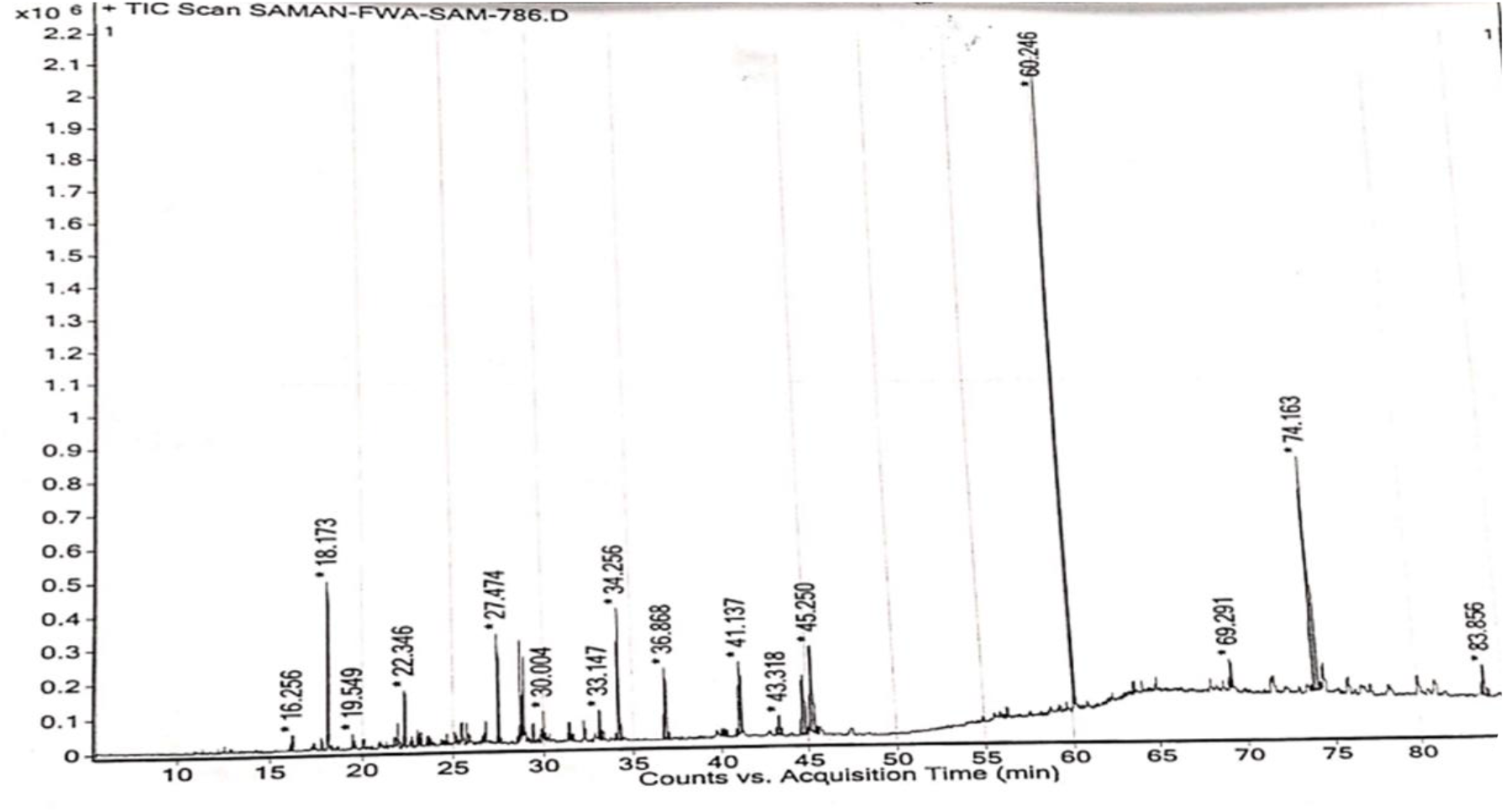
Graph of MS analysis for phytochemicals from the Ethanolic crude extract.

**Figure 8:**
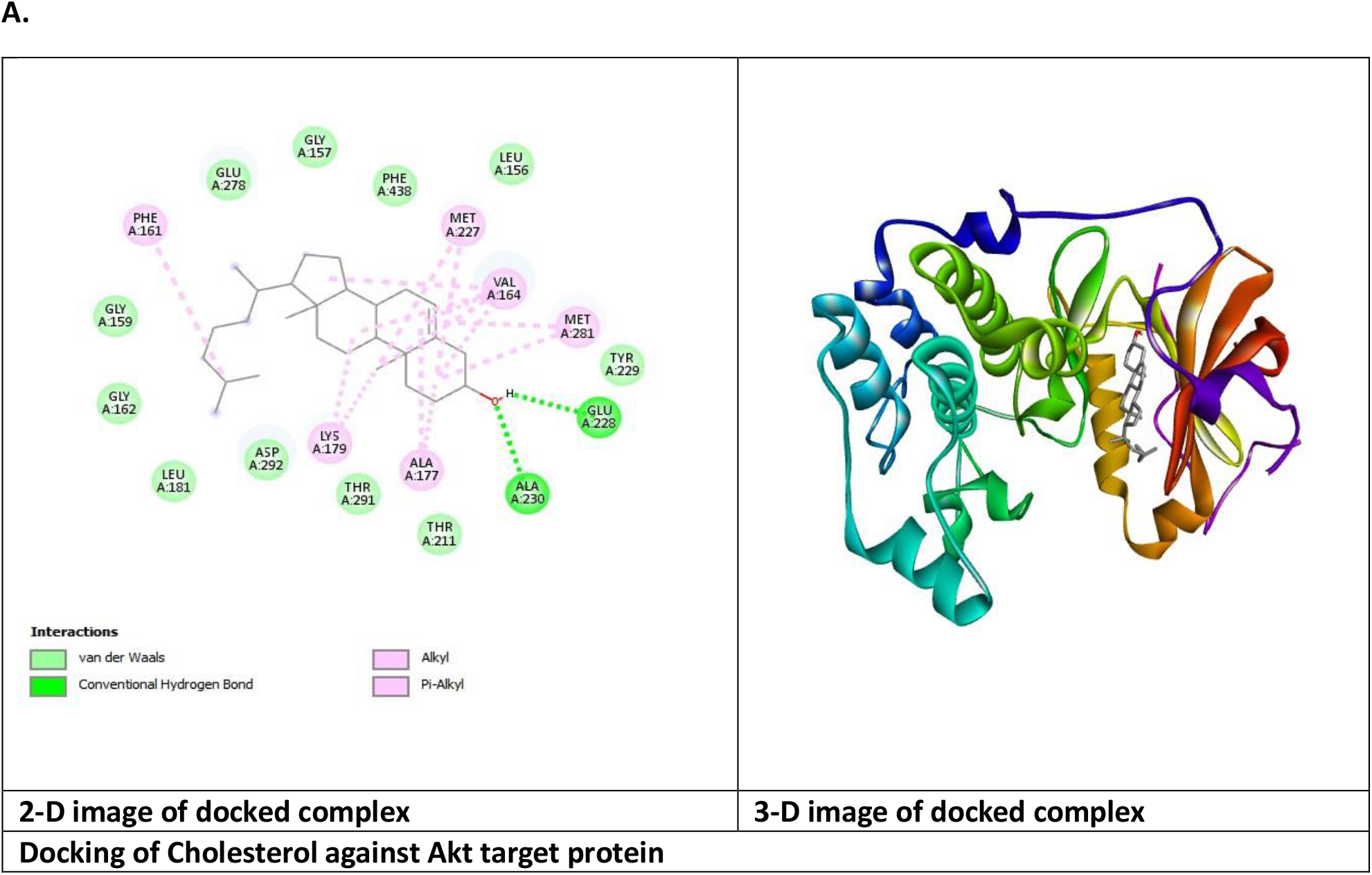
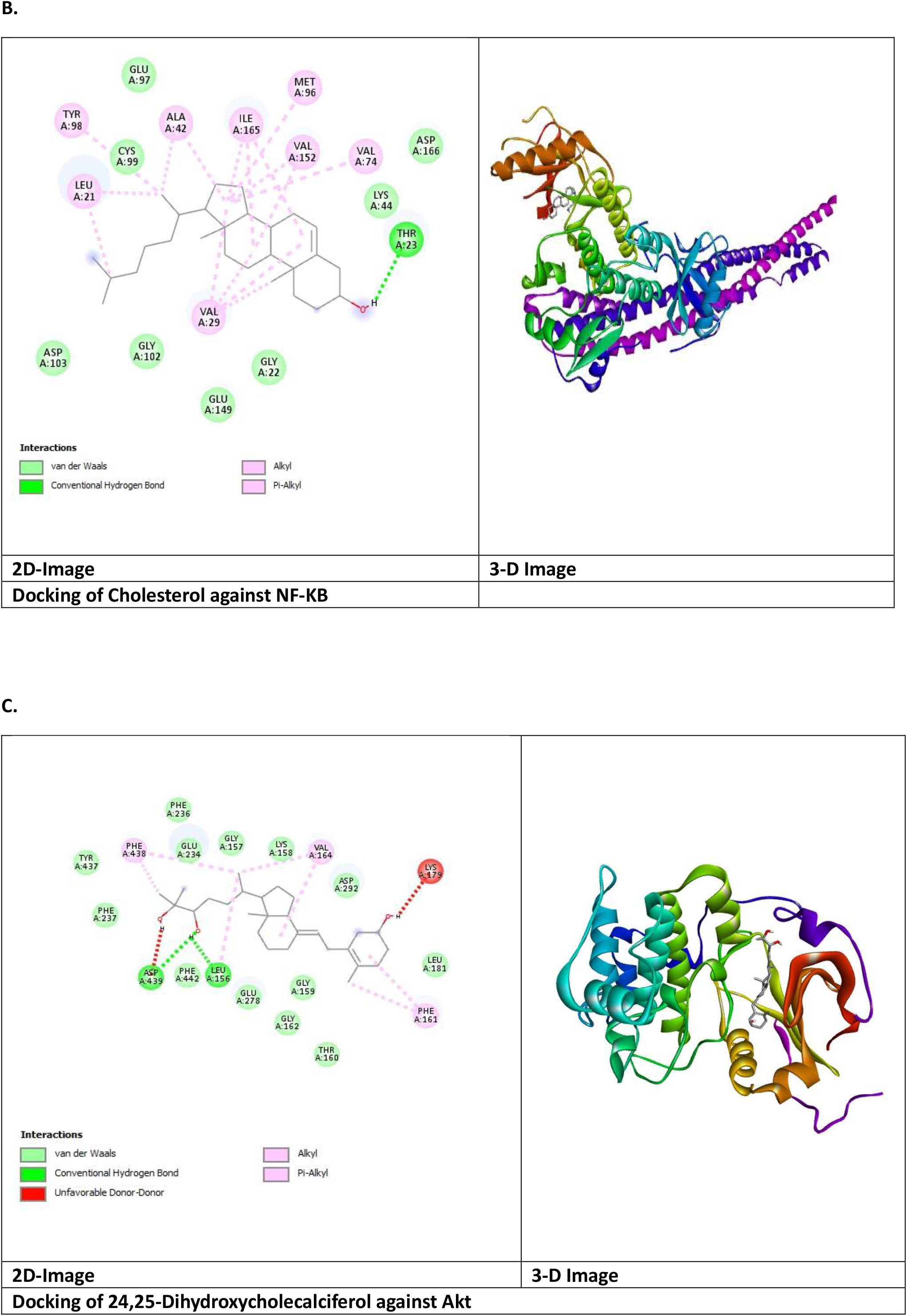
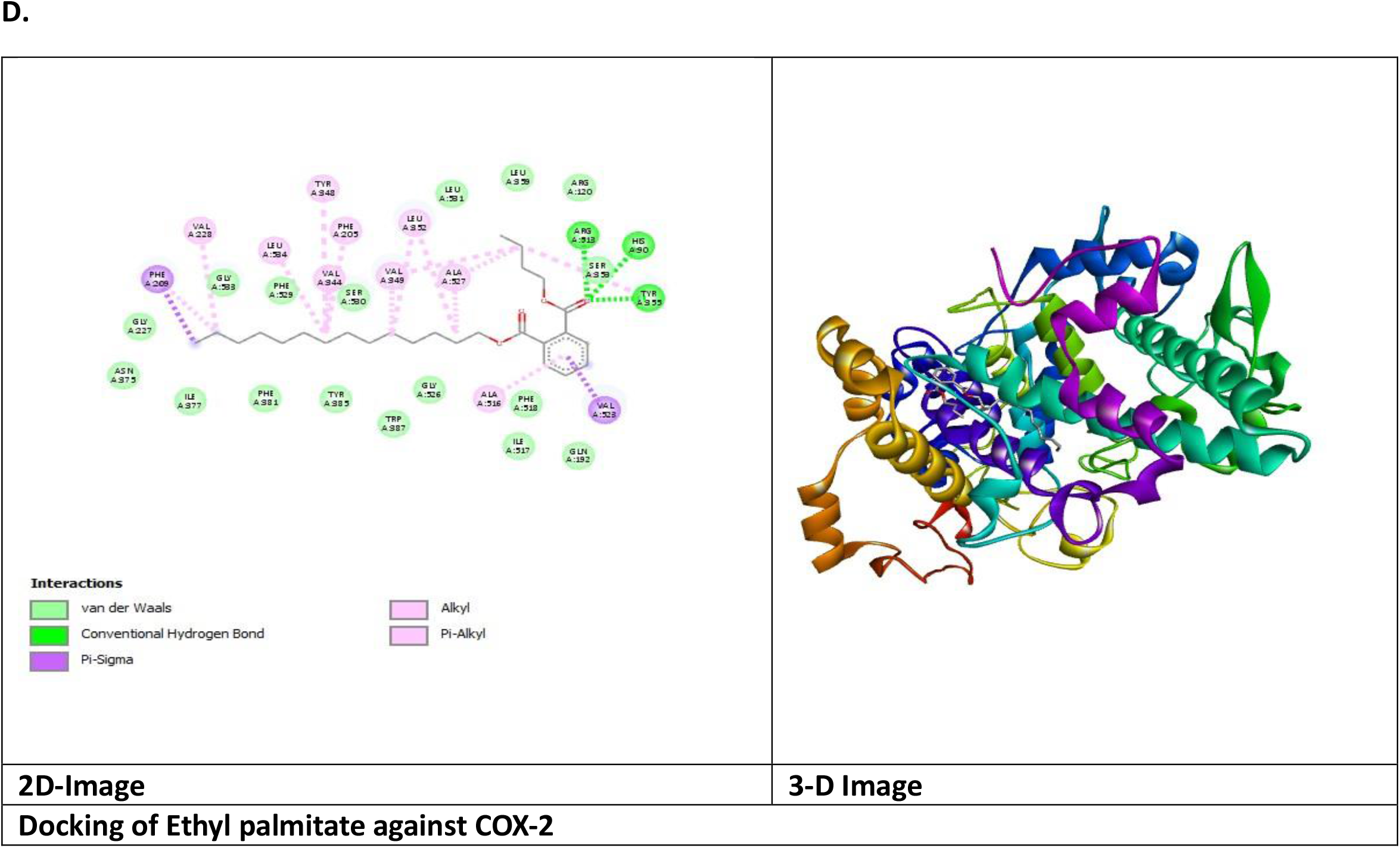
Molecular docking for compounds like A. Cholesterol against Akt target protein, B. Cholesterol against NF-KB24,C. 25-Dihydroxycholecalciferol against Akt. D. Ethyl palmitate against COX-2.

## References

Abbas HAS, Abd El-Karim SS, Ahmed EM, Eweas AF, El-Awdan SA (2016) Synthesis, biological evaluation and molecular docking studies of aromatic sulfonamide derivatives as anti-inflammatory and analgesic agents. Acta Pol Pharm 73(5):1163–1180.

Abid M, Sultana V, Zaidi MJ, Maqbool MA (1997) Nematicidal properties of Stoechospermum marginatum, a seaweed. Pak J Phytopathol 9(2):143–147.

Aftab, S. and Shakoori, A.R., 2019. Low glucose availability alters the expression of genes involved in initial adhesion of human glioblastoma cancer cell line SF767. Journal of Cellular Biochemistry, 120(10), pp.16824–16839.

Ahmad S, Arfan M, Khan AL, Ullah R, Hussain J, Muhammad Z, et al. (2011) Allelopathy of Teucrium royleanum Wall. ex Benth. from Pakistan. J Med Plants Res 5(5):765–772.

Aliyu AB, Musa AM, Sallau MS, Oyewale AO (2009) Proximate composition, mineral elements and anti-nutritional factors of Anisopus mannii NE Br. Trends Appl Sci Res 4(1):68–72.

Amornlerdpison D, Mengumphan K, Thumvijit S, Peerapornpisal Y (2011) Antioxidant and anti-inflammatory activities of freshwater macroalga, Cladophora glomerata Kützing. Thai J Agric Sci 44(5):283–291.

Awad NE, Ibrahim NA, Matloub AA (2009) Phycochemical and cytotoxic activity of some marine algae. Planta Med 75(09).

Canan I, Gündoğdu M, Seday U, Oluk CA, Karaşahin Z, Eroğlu EÇ, et al. (2016) Determination of antioxidant, total phenolic, total carotenoid, lycopene, ascorbic acid, and sugar contents of citrus species and mandarin hybrids. Turk J Agric For 40(6):894–899.

Chung KT, Wong TY, Wei CI, Huang YW, Lin Y (1998) Tannins and human health: A review. Crit Rev Food Sci Nutr 38(6):421–464.

Claudio F, Stendardo B (1965) Experimental and clinical contribution on the use of phycolloid in oncology. Minerva Med 56(85):3617–3622.

Cowan MM (1999) Plant products as antimicrobial agents. Clin Microbiol Rev 12(4):564– 582.

Culioli G, Daoudi M, Ortalo-Magné A, Valls R, Piovetti L (2001) (S)-12-Hydroxygeranylgeraniol-derived diterpenes from the brown alga Bifurcaria bifurcata. Phytochemistry 57(4):529–535.

Dapkevicius A, Venskutonis R, Van Beek TA, Linssen JPH (1998) Antioxidant activity of extracts obtained by different isolation procedures from some aromatic herbs grown in Lithuania. J Sci Food Agric 77(1):140–146.

Fabrowska J, Ibáñez E, Łęska B, Herrero M (2016) Supercritical fluid extraction as a tool to valorize underexploited freshwater green algae. Algal Res 19:237–245.

Findlay JA, Patil AD (1984) Antibacterial constituents of the diatom Navicula delognei. J Nat Prod 47(5):815–818.

Friedman HS, Kerby T, Calvert H (2000) Temozolomide and treatment of malignant glioma. Clin Cancer Res 6(7):2585–2597.

Fujihara M, Iijima N, Yamamoto I, Nagumo T (1984) Purification and chemical and physical characterization of an antitumor polysaccharide from the brown seaweed Sargassum fulvellum. Carbohydr Res 125(1):97–106.

GülçinİSat IG, Beydemir S, Küfrevioglu ÖI (2004) Evaluation of the in vitro antioxidant properties of broccoli extracts (Brassica oleracea L.). Ital J Food Sci 16(1):17–30.

Gupta D (2015) Methods for determination of antioxidant capacity: A review. Int J Pharm Sci Res 6(2):546–566.

Halliwell B, Aruoma OI (1991) DNA damage by oxygen-derived species: Its mechanism and measurement in mammalian systems. FEBS Lett 281(1–2):9–19.

Hanyuda T, Wakana I, Arai S, Miyaji K, Watano Y, Ueda K (2002) Phylogenetic relationships within Cladophorales (Ulvophyceae, Chlorophyta) inferred from 18S rRNA gene sequences. J Phycol 38(3):564–571.

Heil M, Baumann B, Andary C, Linsenmair KE, McKey D (2002) Extraction and quantification of condensed tannins as a measure of plant anti-herbivore defence. Naturwissenschaften 89(11):519–524.

Henkanatte-Gedera SM, Selvaratnam T, Caskan N, Nirmalakhandan N, Van Voorhies W, Lammers PJ (2015) Algal-based single-step treatment of urban wastewaters. Bioresour Technol 189:273–278.

Herrero M, Martín-Álvarez PJ, Señoráns FJ, Cifuentes A, Ibáñez E (2005) Optimization of accelerated solvent extraction of antioxidants from Spirulina platensis. Food Chem 93(3):417–423.

Higgins SN, Malkin SY, Howell ET, Guildford SJ, Campbell L, Hiriart-Baer V, et al. (2008) An ecological review of Cladophora glomerata in the Laurentian Great Lakes. J Phycol 44(4):839–854.

Jeeva S, Marimuthu J, Domettila C, Anantham B, Mahesh M (2012) Preliminary phytochemical studies on selected seaweeds from Gulf of Mannar, India. Asian Pac J Trop Biomed 2(1):30–33.

John DM, Whitton BA, Brook AJ (2002) The Freshwater Algal Flora of the British Isles. Cambridge: Cambridge University Press.

John DM, Rindi F (2015) Filamentous (nonconjugating) and plantlike green algae. In: Wehr JD, Sheath RG, Kociolek JP, editors. Freshwater Algae of North America. 2nd ed. Amsterdam: Elsevier. pp. 375–427.

Kalsoom, A., Altaf, A., Sattar, H., Maqbool, T., Sajjad, M., Jilani, M.I., Shabbir, G. and Aftab, S., 2024. Gene expression and anticancer evaluation of Kigelia africana (Lam.) Benth. Extracts using MDA-MB-231 and MCF-7 cell lines. Plos one, 19(6), p.e0303134.

Kandaswami C, Lee LT, Lee PPH, Hwang JJ, Ke FC, Huang YT, et al. (2005) The antitumor activities of flavonoids. In Vivo 19(5):895–909.

Kang J, Wen Z (2015) Use of microalgae for mitigating ammonia and CO2 emissions from animal production operations. Algal Res 11:204–210.

Koleva II, Van Beek TA, Linssen JPH, De Groot A, Evstatieva LN (2002) Screening of plant extracts for antioxidant activity. Phytochem Anal 13(1):8–17.

Kolodziej H, Kiderlen AF (2005) Antileishmanial activity and immunomodulatory effects of tannins. Phytochemistry 66(17):2056–2071.

Koster JT (1955) The genus Rhizoclonium Kütz. in the Netherlands. Pubbl Stn Zool Napoli 27:335–357.

Kumar KS, Dahms HU, Won EJ, Lee JS, Shin KH (2015) Microalgae: A promising tool for heavy metal remediation. Ecotoxicol Environ Saf 113:329–352.

Laungsuwon R, Chulalaksananukul W (2013) Antioxidant and anticancer activities of freshwater green algae. Maejo Int J Sci Technol 7(2):181–190.

Lee JC, Hou MF, Huang HW, Chang FR, Yeh CC, Tang JY, et al. (2013) Marine algal natural products with antioxidative, anti-inflammatory and anticancer properties. Cancer Cell Int 13(1):55.

Leliaert F, Verbruggen H, Vanormelingen P, Steen F, López-Bautista JM, Zuccarello GC, et al. (2016)DNA-based species delimitation in algae. Eur J Phycol 49(2):179–196.

Louis DN, Ohgaki H, Wiestler OD, Cavenee WK, Burger PC, Jouvet A, et al. (2007) The 2007 WHO classification of tumours of the central nervous system. Acta Neuropathol 114(2):97–109.

Mungmai L, Jiranusornkul S, Peerapornpisal Y, Sirithunyalug B, Leelapornpisid P (2014) Extraction, characterization and biological activities of extracts from freshwater macroalga Rhizoclonium hieroglyphicum. Chiang Mai J Sci 41(1):14–26.

Okwu DE (2001) Evaluation of the chemical composition of indigenous spices and flavouring agents. Glob J Pure Appl Sci 7(3):455–459.

Paya M, Ferrándiz ML, Sanz MJ, Bustos G, Blasco R, Ríos JL, et al. (1993) Study of the antioedema activity of seaweed and sponge extracts. Phytother Res 7(2):159–162.

Rasool M, Iqbal J, Malik A, Ramzan HS, Qureshi MS, Asif M, et al. (2014) Hepatoprotective effects of Silybum marianum and Glycyrrhiza glabra. Evid Based Complement Alternat Med 2014:641597.

Rennie J, Rusting R (1996) Making headway against cancer. Sci Am 275(3):56–59.

Rodríguez-Meizoso I, Jaime L, Santoyo S, Cifuentes A, García-Blairsy Reina G, Señoráns FJ, et al. (2008) Pressurized fluid extraction of bioactive compounds from Phormidium species. J Agric Food Chem 56(10):3517–3523.

Savithramma N, Rao ML, Rukmini K, Devi PS (2011) Antimicrobial activity of silver nanoparticles synthesized using medicinal plants. Int J ChemTech Res 3(3):1394–1402.

Scalbert A (1991) Antimicrobial properties of tannins. Phytochemistry 30(12):3875–3883.

Solanki R, Khanna M, Lal R (2008) Bioactive compounds from marine actinomycetes. Indian J Microbiol 48(4):410–431.

Surayot U, Lee JH, Kanongnuch C, Peerapornpisal Y, Park W, You S (2016) Structural characterization of sulfated arabinans extracted from Cladophora glomerata. Biosci Biotechnol Biochem 80(5):972–982.

Terpinc P, Čeh B, Ulrih NP, Abramovič H (2012) Correlation between antioxidant properties and total phenolic content. Ind Crops Prod 39:210–217.

Varshney A, Singh V (2013) Effects of algal compounds on cancer cell line. J Exp Biol 1:337–352.

Xiao B, Guo J, Liu D, Zhang S (2007) Aloe-emodin induces G2/M arrest in human oral cancer cells. Oral Oncol 43(9):905–910.

Zhang J, Stevens MFG, Bradshaw TD (2012) Temozolomide: Mechanisms of action, repair and resistance. Curr Mol Pharmacol 5(1):102–114.

