## Supplementary data for "Phytochemical profiling and antioxidant potential of freshwater algal extracts from Lahore, Pakistan, with preliminary evaluation of cytotoxic activity"

### S1: Table for GC-MS phytochemical analysis

| GC-MS analysis report | | | | | | | |
|---|---|---|---|---|---|---|---|
| **No of Peaks** | **List of Compounds** | **Molecular Weight** | **Formula** | **Retention Time** | **Area sum %** | **Class of Phytochemical** | **Structure** |
| 1 | Methyl Salicylate | 152 | $C_8H_8O_3$ | 16.256 | 0.2 | Saturated Hydrocarbon | 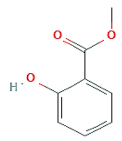 |
| 2 | Thymol | 150 | $C_{10}H_{14}O$ | 18.173 | 2.81 | Volatile oil | 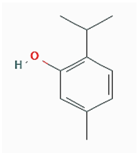 |
| 3 | 3-allyl-6-methoxyphenol | 164 | $C_{10}H_{12}O_2$ | 19.549 | 0.19 | Polyphenol | 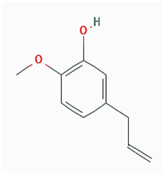 |

| | | | | | | | |
|---|---|---|---|---|---|---|---|
| 4 | 2,4-Di-tert-butyl phenol | 206 | $C_{14}H_{22}O$ | 22.346 | 0.86 | Polyphenol | 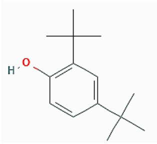 |
| 5 | Actinidiolide, dihydro- | 180 | $C_{11}H_{16}O_2$ | 23.037 | 0.24 | Non-cyclic tetraterpenoid | 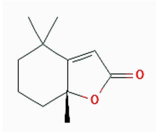 |
| 6 | Elemol | 222 | $C_{15}H_{26}O$ | 23.208 | 0.17 | Sesquiterpenoid | 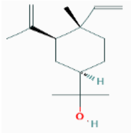 |
| 7 | Ethyl laurate | 228 | $C_{14}H_{28}O_2$ | 23.629 | 0.15 | Saturated Alkane | 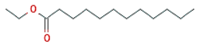 |
| 8 | Myristic acid | 228 | $C_{14}H_{28}O_2$ | 26.776 | 0.55 | Saturated fatty acid | 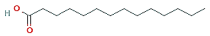 |
| 9 | Ethyl myristate | 256 | $C_{16}H_{32}O_2$ | 27.474 | 2.87 | Saturated Alkane | 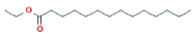 |
| 10 | 3,7,11,15-Tetramethyl-2-hexadecen-1-ol | 296 | $C_{20}H_{40}O$ | 28.718 | 3.04 | Terpenoid alcohol | 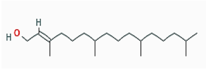 |

| | | | | | | | |
|---|---|---|---|---|---|---|---|
| 1 | Hexahydrofarnesyl acetone | 268 | $C_{18}H_{36}O$ | 28.907 | 2.29 | Ketone | 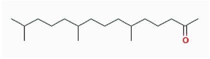 |
| 2 | SAM *1 | 418 | | 30.004 | 0.84 | Alkane ester | |
| 3 | Ethyl palmitate | 284 | $C_{18}H_{36}O_2$ | 31.43 | 0.93 | Alkane ester | 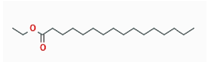 |
| 4 | Arachidonic acid | 304 | $C_{20}H_{32}O_2$ | 33.147 | 1.31 | Carboxylic acid | 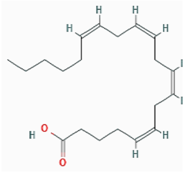 |
| 5 | Methyl oleate | 296 | $C_{19}H_{36}O_2$ | 34.256 | 6.82 | Alkane ester | 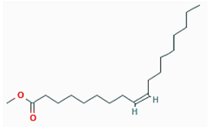 |
| 6 | Phytol | 296 | $C_{20}H_{40}O$ | 36.868 | 4.3 | Saturated Alcohol | 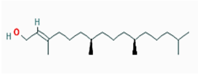 |
| 7 | Dihomo-ɣ-linolenic acid | 306 | $C_{20}H_{34}O_2$ | 40.2 | 0.48 | Carboxylic acid | 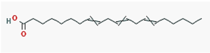 |
| 8 | 9,12-Octadecadienoic acid, ethyl ester | 308 | $C_{20}H_{36}O_2$ | 41.137 | 5.61 | Esters | 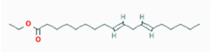 |

| | | | | | | | |
|---|---|---|---|---|---|---|---|
| 9 | Ethyl Oleate | 310 | $C_{20}H_{38}O_2$ | 43.318 | 1.54 | Esters | 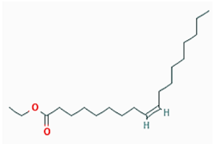 |
| 0 | Monoethylhexyl phthalate | 278 | $C_{16}H_{22}O_4$ | 44.74 | 5.42 | n-alkane | 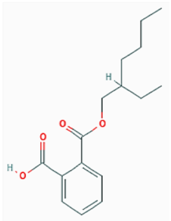 |
| 1 | Octyl phthalate | 390 | $C_{24}H_{38}O_4$ | 45.25 | 7.98 | Esters | 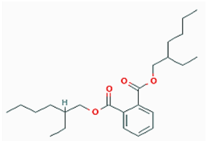 |
| 2 | 3-hydroxycholestan-5-yl acetate | 446 | $C_{29}H_{50}O_3$ | 60.246 | 25.79 | n-alkane | 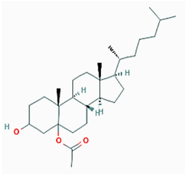 |
| 3 | Cholesterol | 386 | $C_{27}H_{46}O$ | 69.291 | 1.47 | n-alkane | 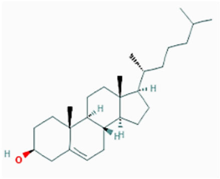 |

| | | | | | | | |
|---|---|---|---|---|---|---|---|
| 4 | ɣ-Sistosterol | 414 | $C_{29}H_{50}O$ | 74.163 | 21.57 | Sterol | 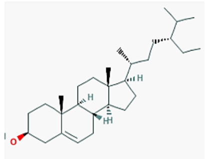 |
| 5 | SAM*2 | 416 | | 83.856 | 2.57 | | |

### S2: Table Docking scores of different compounds against target proteins for crude extract

| Sr. no. | Compound's name | COX-2(PDB ID:5KIR) | Akt (PDB ID:3QKM) | NF-KB (PDB ID:4KIK) |
|---|---|---|---|---|
| 1- | Methyl Salicylate | -5.9 | -5.6 | -6.0 |
| 2- | Thymol | -6.4 | -6.2 | -7.0 |
| 3- | 3-allyl-6-methoxyphenol | -6.6 | -5.8 | -6.3 |
| 4- | 2,4-Di-tert-butyl phenol | -7 | -6.5 | -7.0 |
| 5- | Actinidiolide, dihydro- | -6 | -5.6 | -5.4 |
| 6- | Elemol | -6.9 | -6 | -7.0 |
| 7- | Ethyl laurate | -6.1 | -4.4 | -5.7 |
| 8- | Myristic acid | -6.1 | -4.9 | -5.9 |
| 9- | Ethyl myristate | -6.7 | -4.5 | -5.8 |
| 10- | 3,7,11,15-Tetramethyl-2-hexadecen-1-ol | -7.4 | -5.5 | -6.4 |
| 11- | Hexahydrofarnesyl acetone | -6.6 | -5 | -6.4 |
| 12- | Phthalic acid, butyl tetradecyl ester | -8.1 | -4.7 | -6.4 |
| 13- | **Ethyl palmitate** | **-8.4** | -5.5 | -6.7 |
| 14- | Arachidonic acid | -8.1 | -6.1 | -7.1 |

| | | | | |
|---|---|---|---|---|
| 15- | Methyl oleate | -7.2 | -5.2 | -6.2 |
| 16- | Phytol | -7.3 | -6.2 | -6.6 |
| 17- | 9,12-Octadecadienoic acid, ethyl ester | -7.4 | -4.6 | -6.1 |
| 18- | Ethyl Oleate | -7.1 | -4.6 | -6.3 |
| 19- | Monoethylhexyl phthalate | -7.8 | -5.6 | -7.2 |
| 20- | 3-hydroxycholestan-5-yl acetate | -8.1 | -6 | -7.0 |
| 21- | Octyl phthalate | -7.4 | -5.7 | -7.1 |
| 22- | **Cholesterol** | -7.4 | **-8.6** | **-10.1** |
| 23- | **24,25-Dihydroxycholecalciferol** | -6.5 | **-9.2** | -8.1 |

### S3: Druglikeness (Lipinski's rule of five) of compounds with high binding affinity.

| Sr. no. | Compound's name | Molecular weight<500 | H-Bond donor | H-bond acceptor | LogP<5 | TPSA (20-130Å) | No. of violation (Lipinski's rules) |
|---|---|---|---|---|---|---|---|
| 1- | Cholesterol | 386.65 | 1 | 1 | 4.96 | 20.23Å | 1 vio |
| 2- | 24,25-Dihydroxycholecalciferol | 416.65 | 3 | 3 | 4.38 | 60.69 | 1 vio |
| 3- | Ethyl Palmitate | 284.48 | 0 | 2 | 4.65 | 26.30 | 1 vio |

### S4: ADMET (Pharmacokinetics) of selected compounds

| Property | Model name | Cholesterol | 24,25-Dihydroxycholecalciferol | Ethyl Palmitate |
|---|---|---|---|---|
| Absorption | Water solubility | **-6.917** | **-5.672** | **-7.141** |
| Absorption | Caco2 permeability | **1.214** | **1.112** | **1.596** |
| Absorption | Intestinal absorption (human) | **93.723** | **92.259** | **91.916** |

| | | | | |
|---|---|---|---|---|
| Absorption | Skin Permeability | **-2.841** | **-3.013** | **-2.682** |
| Absorption | P-glycoprotein substrate | **No** | **Yes** | No |
| Absorption | P-glycoprotein I inhibitor | Yes | **Yes** | No |
| Absorption | P-glycoprotein II inhibitor | Yes | **Yes** | No |
| Distribution | BBB permeability | **0.763** | **-0.312** | **0.759** |
| Distribution | CNS permeability | **-1.703** | **-1.879** | **-1.777** |
| Metabolism | CYP2D6 substrate | **No** | **No** | No |
| Metabolism | CYP3A4 substrate | Yes | Yes | Yes |
| Metabolism | CYP1A2 inhibitior | No | No | Yes |
| Metabolism | CYP2C19 inhibitior | No | No | No |
| Metabolism | CYP2C9 inhibitior | No | No | No |
| Metabolism | CYP2D6 inhibitior | No | No | No |
| Metabolism | CYP3A4 inhibitior | No | | No |
| Excretion | Total Clearance | **0.589** | **-1.733** | **1.912** |
| Excretion | Renal OCT2 substrate | **No** | **No** | No |
| Toxicity | AMES toxicity | **No** | No | No |
| Toxicity | Max. tolerated dose (human) | **-0.58** | **-1.733** | **0.199** |
| Toxicity | hERG I inhibitor | **No** | No | No |

| | | | | |
|---|---|---|---|---|
| Toxicity | hERG II inhibitor | **Yes** | No | **No** |
| Toxicity | Oral Rat Acute Toxicity (LD50) | **2.1** | **4.697** | **1.664** |
| Toxicity | Oral Rat Chronic Toxicity (LOAEL) | **0.967** | 2.06 | **3.019** |
| Toxicity | Hepatotoxicity | No | No | **No** |
| Toxicity | Skin Sensitisation | No | **No** | **Yes** |
| Toxicity | *T.Pyriformis* toxicity | **0.626** | **0.592** | **1.837** |
| Toxicity | Minnow toxicity | **-1.715** | **1.677** | **-1.538** |
